# Prediction of Optimal Time of Day for Salmeterol Based on Circadian Phases of Target Proteins in the Target Tissues

**DOI:** 10.1101/2025.09.24.673459

**Authors:** Sibel Cal-Kayitmazbatir

**Affiliations:** Department of Musculoskeletal Science and Ageing, University of Liverpool

## Abstract

Circadian rhythms are conserved across a wide range of organisms, including cyanobacteria, fungi, insects, and mammals. Approximately 50% of gene transcription, 8-38% of protein translation, and 13-22% of metabolites in mice have been shown to cycle with 24-hour rhythmicity. Approximately 75% of the current medications on the market target circadian-controlled pathways. However, current drug discovery and improvement studies are far from investigating the effect of time of day. In the current study, the optimal time of day for treating Chronic Obstructive Pulmonary Disease (COPD) with salmeterol is shown based on the circadian profiles of the target proteins in the target tissues. This calculation strategy aims to provide timing guidance for the drugs targeting rhythmic pathways or being metabolised by them. The current study also reveals different predicted optimal times of day for male and female mice, highlighting the importance of gender variability in circadian studies. The approach proposed in this research could be beneficial in studies that consider the circadian timing of medications.

## Introduction

Circadian rhythms are tightly controlled at transcriptional, translational and post-translational levels [1, 2]. Even though there are different levels of internal controls, body rhythms are also affected by external time cues (zeitgebers), which is both a pro and a con [3]. External regulation is a pro because it means clocks can be manipulated, and a con because modern life has numerous time-resetting cues that disrupt the body’s clock.

Circadian rhythms are interlocked with many diseases. While in some cases a disrupted body clock is a result of a disease, it can also be the cause [4]. It is essential to note that even if clock disruption is a result of a disease, improving body clocks can enhance the well-being of individuals and improve disease outcomes [5].

Circadian medicine aims to utilise the power of circadian rhythms and improve diagnosis and treatment strategies. However, only 6.1% of the highly cited papers include time-of-day information, which is significantly lower compared to the pathways related to circadian rhythms [6]. Circadian medicine has three approaches: (1) detecting the clock through biomarkers of blood/skin or wearable devices; (2) exploiting the clock by using circadian rhythms to determine the optimal time of day for drug administration; (3) targeting the clock when the body’s rhythm is disrupted (Figure 1).

**Figure 1:**
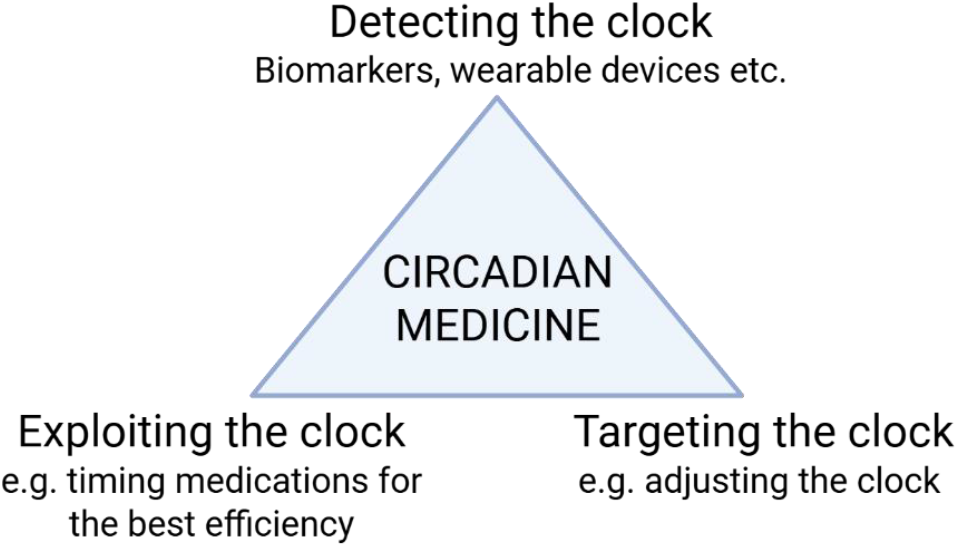
Three main approaches of circadian medicine. Figure adapted from Achim Kramer et. al., 2022, PLoS Biology

Female inclusion in the circadian studies is limited to 18% in mice [7] and 33.9% in humans [8]. However, the distribution of diseases is usually equal for both genders. It is time to incorporate gender variables more into circadian rhythm studies.

Chronic Obstructive Pulmonary Disease (COPD) is a lung condition that causes chronic bronchitis and emphysema in the lung tissue. Chronic bronchitis causes inflammation and excessive mucus production, leading to coughing and narrowed airways. Emphysema causes enlarged alveoli, resulting in shortness of breath [9]. COPD is the 4^th^ leading cause of death worldwide, and every year, 1.2 million people in England are diagnosed with it. COPD has been shown to disrupt lung circadian rhythms and causes muscle wasting, a condition that causes muscle tissue breakdown and muscle weakness. Circadian rhythms are disrupted in lung airway inflammation [10].

While there are many studies investigating the effects of salmeterol on lung inflammation in humans [11] and in mice [12, 13], daily timing studies were limited. Salmeterol was shown to have an overnight protection effect when administered towards the end of the activity phase in a clinical trial [14].

One of the biggest challenges of circadian medicine is the lack of generalisable approaches. Here, by predicting the optimal time of day for drugs using healthy mouse tissues and the circadian phases of drug target proteins, we can create a death worldwide, and every year, 1.2 million people in England are diagnosed with it. COPD has been shown to disrupt lung circadian rhythms and causes muscle wasting, a condition that causes muscle tissue breakdown and muscle weakness. Circadian rhythms are disrupted in lung airway inflammation [10]. method to apply the results of this approach to human studies.

I hypothesize that;

Optimal time of day for drug administration can be calculated based on the protein rhythmicity profiles of the target proteins and drug metabolising proteins in the target tissues, and the protein of the target in the side effect-related organ, if it exists. When this is being calculated, the drug’s administration route and its diffusion time to reach the target tissues should be considered (Figure 2).

**Figure 2:**
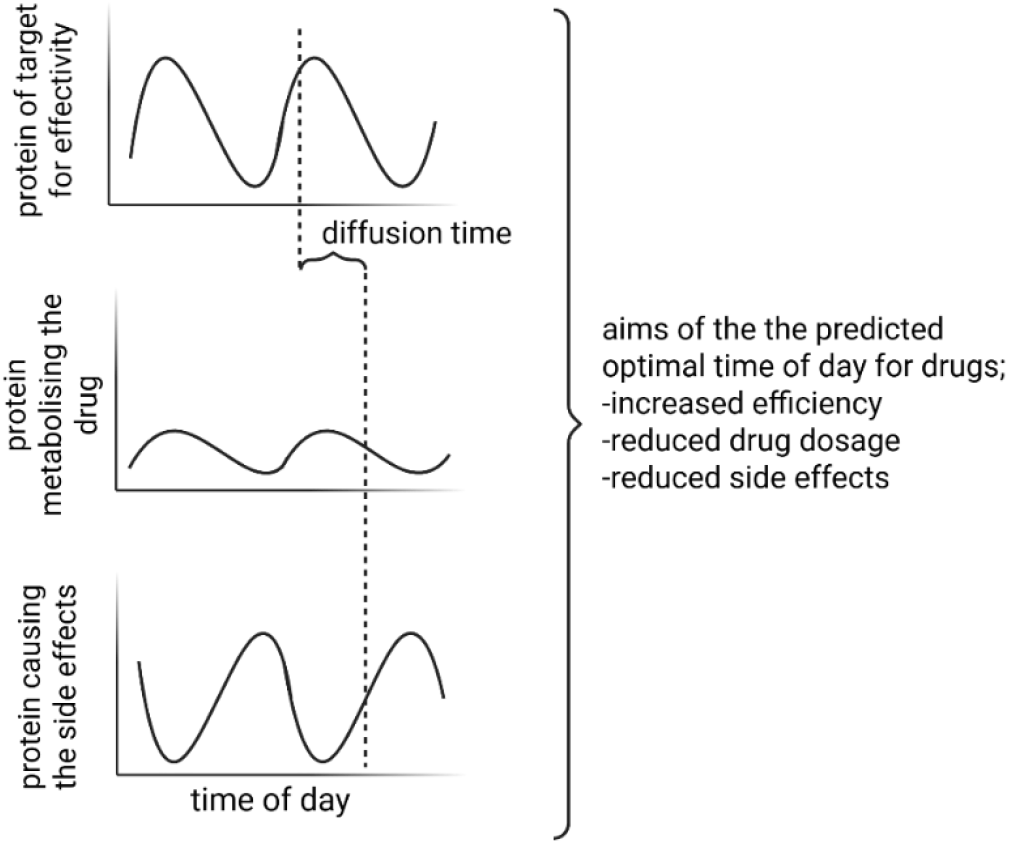
Proposed method of predicting optimal time of day for drug administration

## Materials and Methods

### Prediction of optimal time of day for drugs

Circadian medicine studies typically involve testing different times of day in human or mouse studies to determine the effects of time on treatment outcomes. Different times usually involve AM and PM dosages. While this is the currently accepted approach, it is time-consuming and resource-intensive and doesn’t allow for precise timing. More accurate timing trials must follow the initial broad timing study (AM vs PM). The proposed prediction can guide researchers in targeting potentially optimal times and can show if gender affects the phases of the drug target proteins.

The prediction involves six main steps:

1. Determination of the drug,
2. Determination of the target proteins and target tissues,
3. Determination of the circadian profiles of the target proteins in the target tissues,
4. Literature search for the drug’s ADME (absorption, distribution, metabolism, excretion) properties,
5. Circadian analysis of the protein profiles through the meta2d R package
6. Calculation of the predicted time of day to target the highest target protein level in the target tissue for the highest efficiency, and the lowest level of drug metabolising protein, and the lowest level of side effect-causing protein after diffusion time, as shown in Figure 2

### Animals

3-month-old healthy male and female C57BL/6 mouse tissues were used for the study. Mice were kept under 12:12 light:dark cycle where ZT0 corresponds to the time lights come on. Mice were maintained in the Biomedical Sciences Unit of the University of Liverpool and euthanised with cervical dislocation under Home Office Regulations. Collected tissues were snap-frozen and processed later.

### Determining the circadian profiles of drug target proteins

Protein samples were prepared in RIPA buffer. Bio-Rad SDS-PAGE electrophoresis system and western blot method were used to determine the protein expression phenotypes of ADRB2 (Proteintech 29864-1-AP), CYP3A4 (abcam ab3572), and PER2 (STJ115134 proteins.Calculation of circadian parameters of the western blot results.Meta2d R package was used to calculate circadian parameters of the protein expression results.

## Results

### Proof of concept application of the optimal time of day prediction to chronic obstructive pulmonary disease (COPD) treatment with salmeterol

Salmeterol is one of the bronchodilators used to treat COPD and has been shown to have the potential to alleviate muscle wasting [15]. Salmeterol is an inhaled drug which immediately reaches the lungs when administered, and it takes one to two hours for it to reach the liver to be metabolised and the muscle to help with the wasting phenotype. To determine the circadian profiles of salmeterol target proteins in its target organs, the liver, lung and quadriceps muscles of 3-month-old male and female healthy C57BL/6 mice were collected and processed for western blot. Supplementary Figure 1 shows the Ponceau S staining images for all the membranes used.

To determine if there is a relationship between the phase of PER2 in the liver, lung, and quadriceps muscle, and to check for correlations, we next checked PER2 levels in the same samples and analysed the quantification data. Figure 4 shows PER2 western blot every six hours for a day in female (top) and male (bottom) liver, lung, and quadriceps muscle tissues and their quantifications. While the liver CYP3A4 phase correlated with the PER2 phase in female samples, there was an 8-hour shift between the male liver PER2 and CYP3A4 phases. On the other hand, in males, lung ADRB2 phase correlated with lung PER2 phase, which was 11 hours shifted in females. When the amplitude levels are compared in Table 2, opposite to salmeterol target proteins, PER2 amplitudes in male lung and quadriceps samples were higher than those of females.

**Table 1:** Phases of salmeterol target proteins calculated by meta2d R package. The data indicate that F(female) and M(male) protein phases and amplitudes of the rhythmicity are different, which should be considered when predicting the optimal time of day of salmeterol for COPD treatment.

|  | phase |  | amplitude |  |
| --- | --- | --- | --- | --- |
|  | F | M | F | M |
| lung-ADRB2 | 22.07 | 18.79 | 0.19 | 0.107 |
| quad-ADRB2 | 0.33 | -0.55 | 0.16 | 0.108 |
| liver-CYP3A4 | 13.21 | 21.53 | 0.06 | 0.133 |

**Table 2:** PER2 phases and amplitudes in lung, quadriceps and liver in male and female mice calculated by meta2d R package. The data indicate that the PER2 phase is delayed in the lung and quadriceps, while it is advanced in the liver in male mice compared to females.

|  | phase |  | amplitude |  |
| --- | --- | --- | --- | --- |
|  | F | M | F | M |
| lung-PER2 | 9.68 | 19.96 | 0.16 | 0.26 |
| quad-PER2 | 8.03 | 19.78 | 0.06 | 0.33 |
| liver-PER2 | 13.43 | 4.67 | 0.12 | 0.06 |

While most of the data from research is either represented in manuscripts or deposited in archives where data scientists can use interactively, data from the study is implemented in an R Shiny code for better utilisation of the data. Although it is not available online yet due to the costs of the step, a screenshot of locally run code can be seen in Figure 5. In future studies, the application will be made available online.

In Figures 3 and 4, the levels of proteins were calculated relative to ZT2. To show whether there is a difference between the levels of proteins at ZT2 in male and female samples, I analysed these samples with a western blot on the same membrane. Even with a limited sample size, lung ADRB2 and liver PER2 levels were significantly lower in the males (Figure 6).

**Figure 3:**
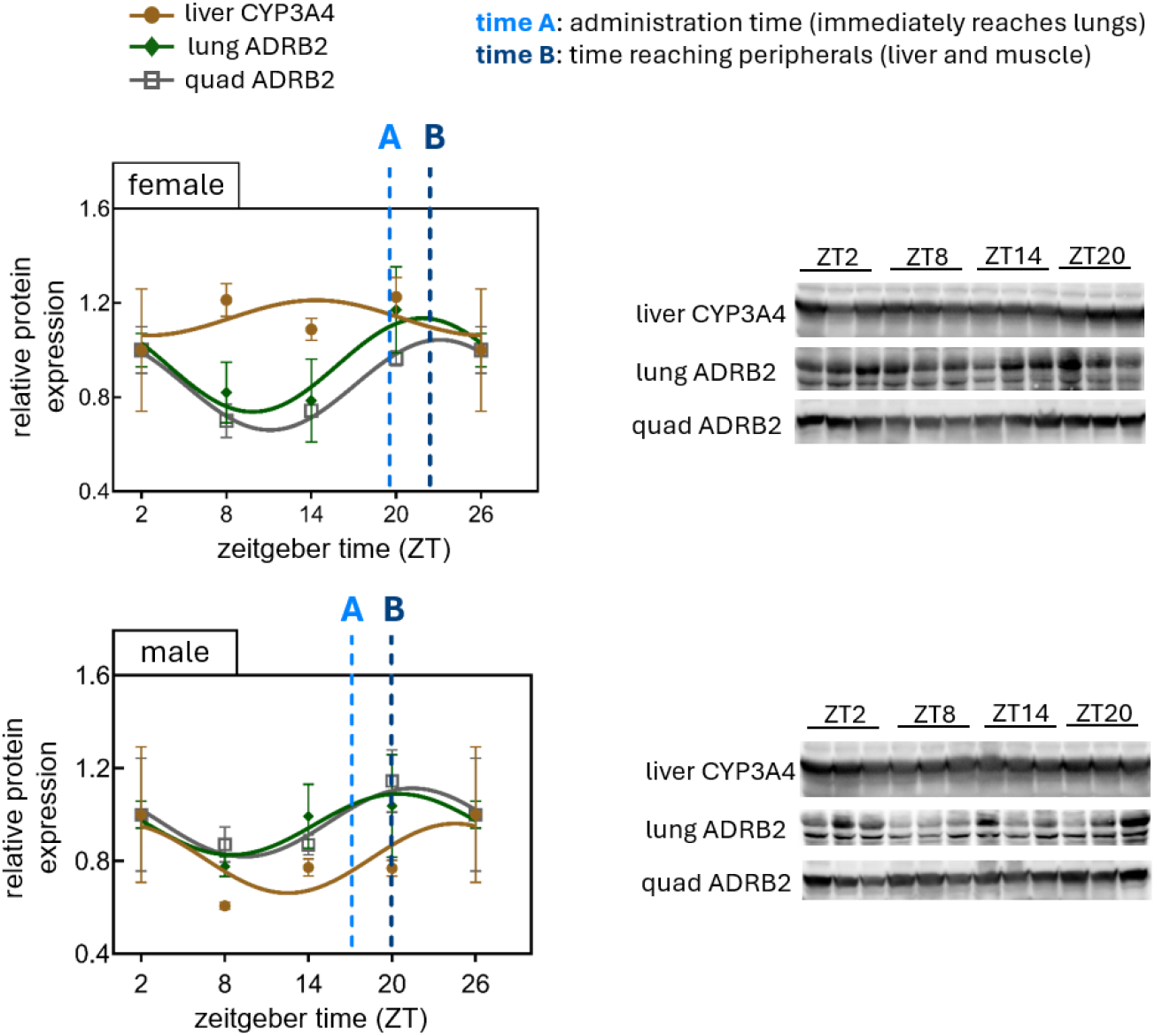
Salmeterol target proteins over time at three different tissues in female (top) and male (bottom) mice. Tissues from male and female C57BL/6 mice were collected every 6 hours and processed for protein extraction and western blot analysis for the indicated proteins. n=3 biological replicates. All target protein levels were normalised to total protein amounts in each well, calculated from Ponceau S staining images (Supplementary Figure 1).

**Figure 4:**
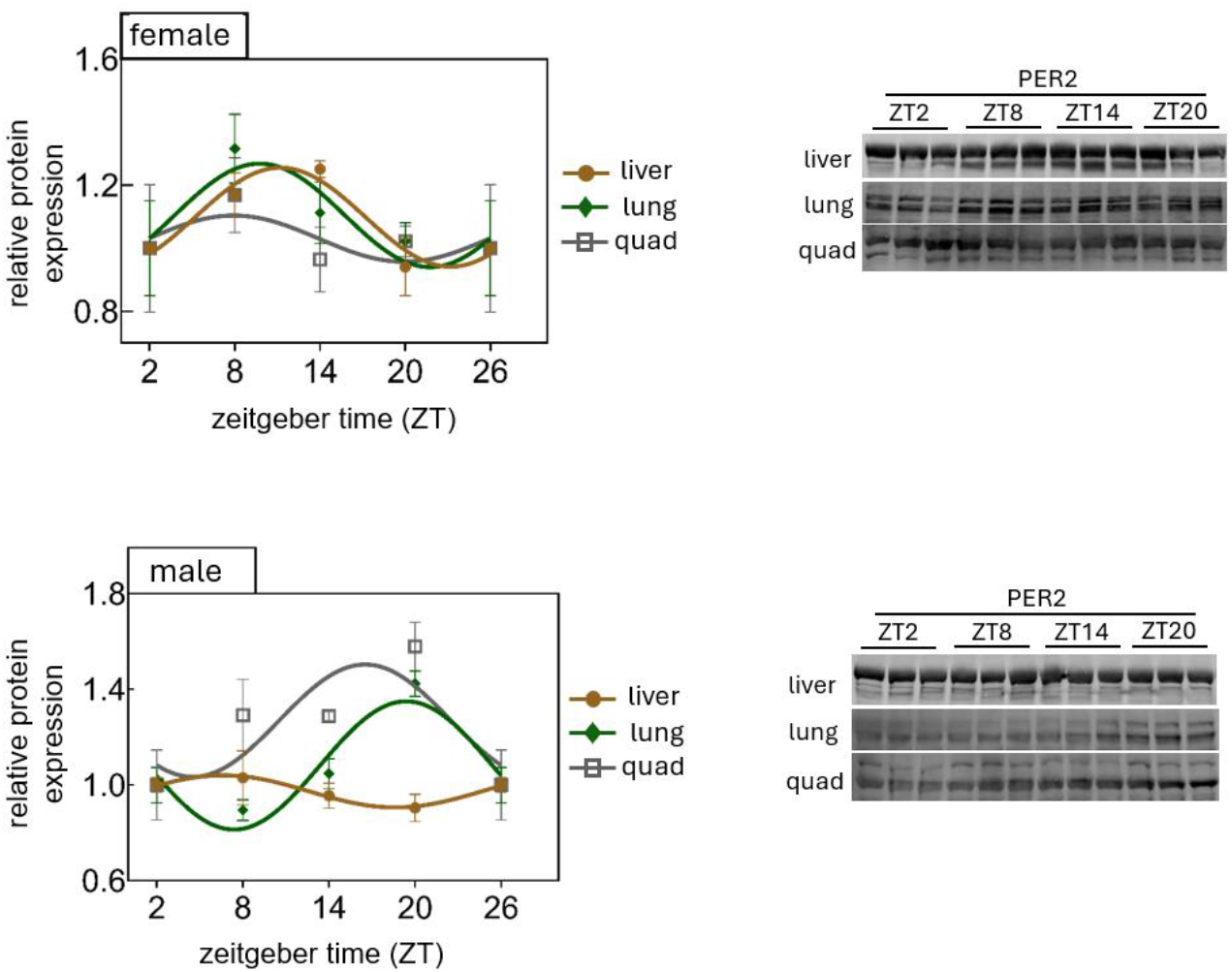
Core circadian clock protein PER2 circadian phases in liver, lung, and quadriceps of male and female C57BL/6 mice. n=3 biological replicates. All target protein levels were normalised to total protein amounts in each well, calculated from Ponceau S staining images.

**Figure 5:**
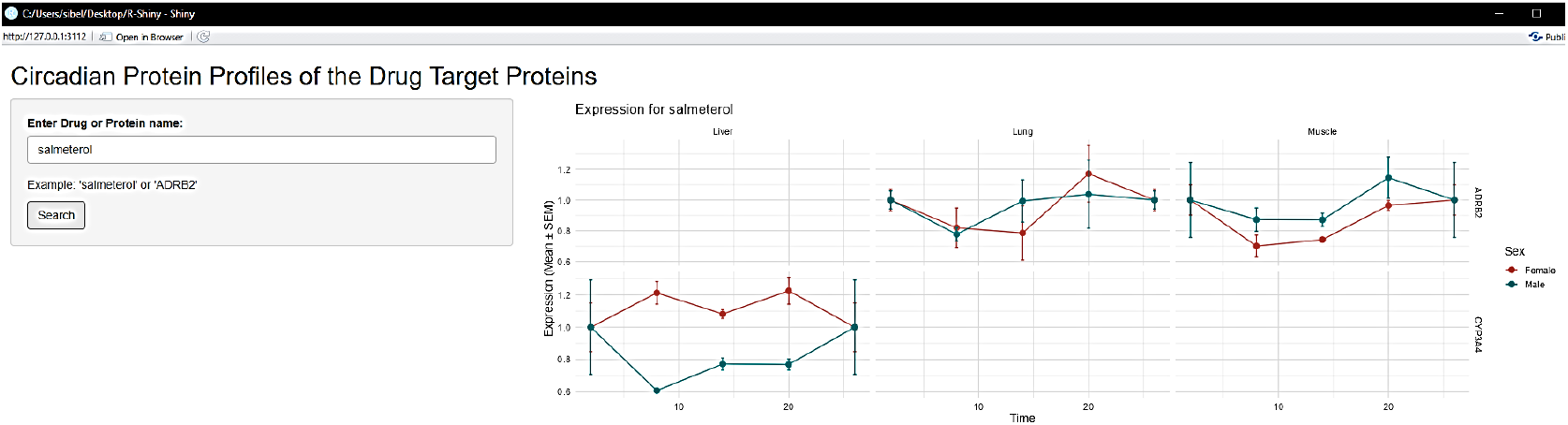
Screenshot of the locally run code for interactively representing circadian protein profiles of the drug target proteins.

**Figure 6:**
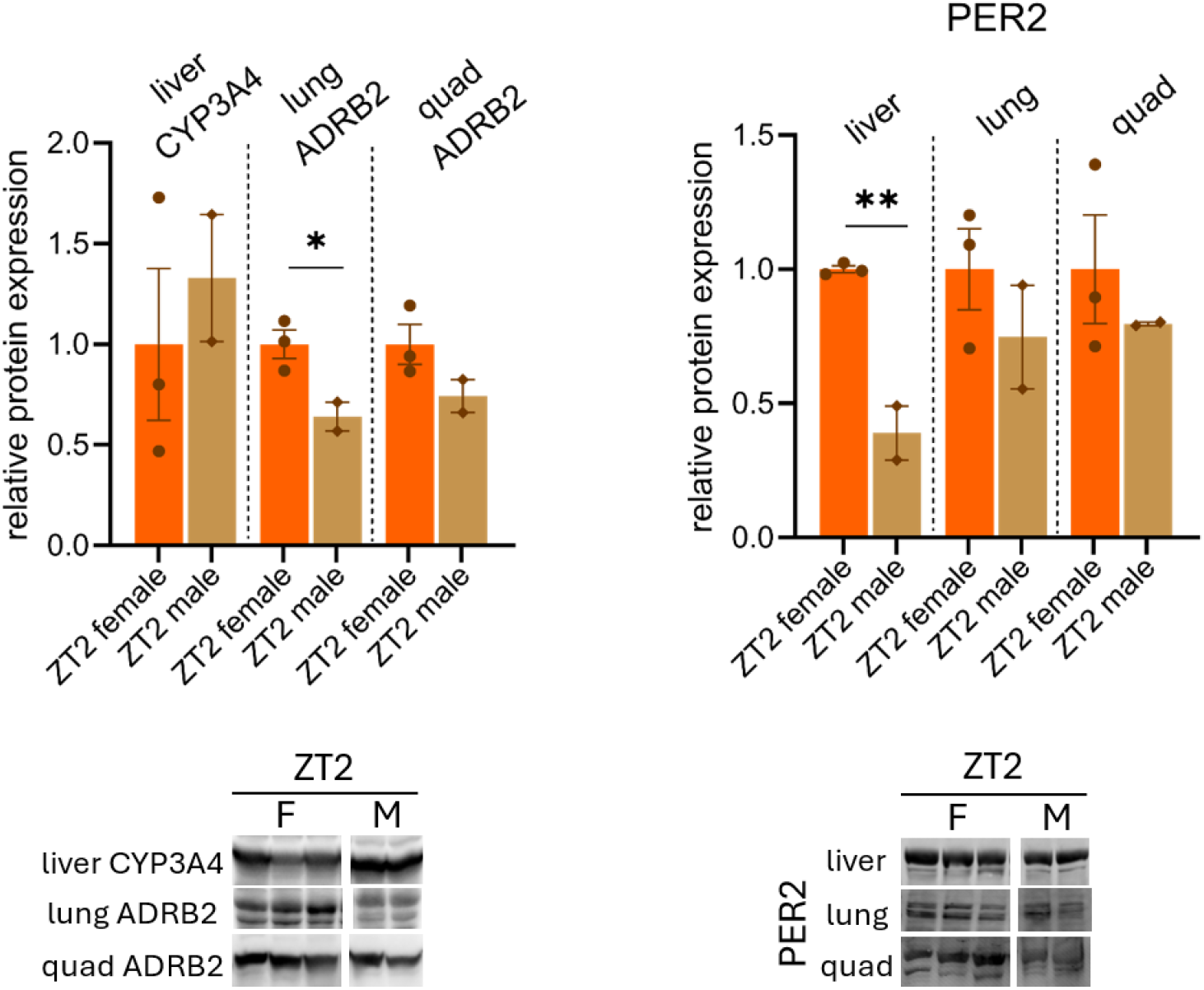
ZT2 protein levels of salmeterol target proteins and PER2 in liver, lung, and quadriceps of male and female mice. n=3 biological replicates for female, n=2 biological replicates for male samples. Welch's t-test was applied to analyse the significance.

Thus, this study highlights a method to predict the optimal time of day for salmeterol using circadian protein phases of the target proteins in different tissues and shows the importance of male and female differences when salmeterol is processed in the body.

To calculate the predicted optimal time in the day,

Find the time of maximum P_target_= t_peak_

As we know the diffusion time for salmeterol, we find the minimum of the score below at two hours after the t_peak_. At that delayed time, t_peak_ +2, computed the score:

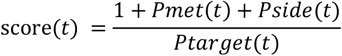

## Discussion

The reason salmeterol was chosen for the study was to verify the efficiency of the prediction and discuss its potential to be the bridge between mice and humans. Although limited, there are some COPD treatment timing clinical studies, and the predicted optimal time of day for salmeterol aligns with the clinical study data [14].

It should be noted that determining one circadian rhythm protein is not enough for understanding any correlations between the clock and salmeterol target proteins for different genders. Future studies should include more circadian rhythm proteins to clarify any relationship between the drug target protein phases.

Drug timing studies are usually performed in diseased mice, whereas I suggest a different method here, where we can predict the optimal time of day for a drug in healthy mice. This approach is a starting point for future studies, which can include determining if the clock is changed in the diseased model, fixing the clock with clock modulators, and applying the predicted optimal time of day for the treatment.

This project was supported by the MAS Research Support Budget 2025 by the Musculoskeletal Science and Ageing Department of the University of Liverpool.

